# Keratinocyte FABP5 mediates saturated fat-induced MAPK/CCL20 signaling in psoriatic inflammation

**DOI:** 10.64898/2026.10.07.757340

**Authors:** Jianyu Yu, Jiaqing Hao, Anthony Avellino, Xingshan Jiang, Xiaochun Han, Ali Jabbari, Bing Li

## Abstract

Saturated fats in high-fat diets (HFDs) have been associated with increased risk and severity of psoriasis. However, the underlying cellular and molecular mechanisms remain to be further defined. Here, using a unique mouse strain with keratinocyte-specific deletion of fatty acid-binding protein 5 (FABP5) (Fabp5^f/f^Krt6a-Cre), we reported that keratinocyte FABP5 deletion significantly attenuated saturated HFD-exacerbated psoriatic symptoms. Immunological analyses demonstrated that FABP5 specific deletion in keratinocytes selectively reduced skin TCR γ δ ^dim^ cell accumulation and IL-17A production. Further skin bulk-RNA and single-cell RNA sequence analyses showed a strong association of keratinocyte FABP5 with increased activation of mitogen-activated protein kinase (MAPK) pathways under saturated HFD-induced obesity. Mechanistically, saturated fatty-acids (SFAs) induce FABP5-dependent MAPK activation and CCL20 production in keratinocytes, promoting the recruitment and activation of IL-17-producing γδ T cells in the skin. Together, these findings suggest a previously undefined FABP5/MAPK/CCL20 signaling axis that links saturated fats to exacerbated psoriatic inflammation.

## Introduction

Driven by a complex combination of genetic susceptibility and environmental factors, psoriasis is an immune-mediated chronic skin inflammation, affecting approximately 125 million people in the world^1, 2^. Accumulating evidence from both clinical studies and experimental models has demonstrated that the crosstalk between keratinocytes and multiple immune cell subsets, particularly through IL-23/IL-17/IL-22 axis, plays a central role in psoriasis pathogenesis^3^. Dietary-induced obesity (DIO) is an important risk factor for psoriasis^4, 5^. Emerging evidence suggests that dietary fat composition may influence psoriatic inflammation independently of adiposity^4, 6^. In particular, saturated fatty acids (SFAs) are the major type of dietary fat relevant to increased psoriasis risk and disease severity by amplifying multiple pro-inflammatory pathways in the skin^7^. However, cellular and molecular mechanisms by which dietary SFAs exacerbate psoriatic inflammation remain incompletely understood.

Keratinocytes, the predominant cell type in epidermis, play a pivotal role in the psoriatic inflammatory circuit by producing cytokines/chemokines that promote recruitment and activation of downstream immune cells, including neutrophils and IL-17-producing T cells^8, 9^. Although skin-resident immune cells, particularly innate immune cells, have been extensively studied for their responses to lipid stimulus, particularly SFAs^10, 11^, emerging evidence highlights keratinocytes as active sensors of metabolic perturbations that can translate metabolic cues into proinflammatory signaling^12, 13^. However, how keratinocytes directly sense elevated SFAs in obesity and integrate metabolic and inflammatory pathways in psoriatic pathogenesis remains poorly understood.

Epidermal fatty acid-binding protein (E-FABP or FABP5) is one of the nine members of FABP family and is abundantly expressed in skin epidermis, facilitating hydrophobic fatty acid intracellular trafficking and utilization^14^. Accumulating evidence from both human and animal studies has established an important role for FABP5 in psoriasis pathogenesis ^15–17^. Our previous study demonstrates that FABP5 in skin macrophages mediates SFA-related psoriatic inflammation^18, 19^. Beyond immune cells, research from our group reveals that FABP5 expression in keratinocytes promotes psoriatic inflammation by interacting with valosin-containing protein (VCP) to enhance NF-κB signaling under non-obese conditions^9, 20^. Given FABP5’s dual functions in intracellular lipid handling and immunoregulation, we hypothesize that FABP5 expression in keratinocytes serves as a molecular link between dietary SFAs and inflammatory responses in obesity-related psoriasis.

In this study, using a keratinocyte-specific Fabp5 conditional knockout mouse model, we demonstrated that keratinocyte FABP5 contributes to saturated high-fat diet (HFD)-exacerbated psoriasis through a mitogen-activated protein kinase (MAPK)/CCL20/ signaling axis that links dietary SFA to dysregulated cutaneous immune responses and exacerbated psoriatic inflammation.

## Materials and Methods

### Mice and saturated high-fat diet (HFD)-related psoriasis

Fabp5^f/f^ mice were bred and housed in the animal facility at the University of Iowa. Krt6A-Cre mice of C57BL/6 background were purchased from Jackson Laboratory. Fabp5^f/f^ Krt6A-Cre mice were generated as previously described ^9^. Male and female mice were weaned at 3–4 weeks of age and randomly assigned to different groups for special diet feeding, which included a low-fat diet (LFD, 5% fat from soybean) and cocoa butter high-fat diet (HFD, 45% fat). Mouse body weight was monitored every three weeks, and body fat deposition was measured via NMR scanning after six months on special diets.

Following five months of dietary treatment, mice were used for psoriasis induction via imiquimod (IMQ) treatment with Vaseline as vehicle-cream control, as previously described [cite]. Briefly, hair was removed from the mice’s back skin using 200mg of Nair (a commercial facial removal cream), and the mice were monitored for another 48 h. The hair-removed mice were then randomly treated with 4 mg of IMQ cream (Perrigo, containing 5% IMQ) for four consecutive days. Disease severity was scored based on erythema, induration, and desquamation as previously described^17^. The cumulative score was calculated as the sum of these three parameters. At the endpoint of the IMQ treatment, mice were sacrificed for further immunophenotypic analysis using flow cytometry in various tissues.

### Single-cell suspension preparation

The preparation of single-cell suspension from the skin epidermis and dermis was conducted following our previous work^17^. Briefly, psoriatic skin lesions from mice were removed. The removed skin was placed with the epidermal side down on the lid of the 60mm culture dish containing 6 mL of cold PBS. Subcutaneous fat was scraped off from the skin using a tweezer and the blunt end of a scalpel. The remaining skin tissue was cut into 0.5 × 1 cm pieces and incubated with 6 mL of 0.25% Trypsin-EDTA solution for 1.5 h at 32°C with gentle shaking at 20 rpm. After the incubation, the epidermal and dermal layers were separated with the blunt end of a scalpel. Epidermal single cells were obtained by thoroughly dispersing the epidermis tissue with a pipette until all the fragments passed freely through the tip of a 1mL pipette. The cell suspension was then filtered through a 40 μm cell strainer. The dermis tissue was minced and further digested in the tri-enzyme skin-digesting buffer containing 0.2 mg/mL hyaluronidase, 0.02 mg/mL DNase-1, and 0.5mg/mL collagenase IV in RPMI-1640 medium supplemented with 5% fetal bovine serum, for 45 min at 37°C with gentle shaking 20 rpm. Following the digestion, the dermis tissue was transferred into a 50mL centrifuge tube and vortexed for at least 20 s. Dermal single cells were then obtained by a two-step filtration process: first through a 105um mesh filter cloth, followed by a 40 μm cell strainer.

For single-cell suspension from draining lymph nodes (dLNs), both lymph nodes from both sides are collected with removing adherent fats and kept in cold PBS of the 60mm tissue culture dish. dLNs are ground in cold PBS to release single cells. Cell suspension was filtered through a 40μm cell stainer into a 15mL centrifuge tube. Cells were then washed twice with cold PBS, pelleted, and resuspend in cold PBS.

For splenic single-cells, mouse spleen was collected and ground to release single cells. Cells were filtered through a 40μm cell stainer into a 15mL centrifuge tube, and pelleted. Cells were then processed red blood cell lysis by resuspending in 3mL RBC lysis buffer (Tonbo Biosciences) and incubated for 3 min at room temperature. Cells were then washed by cold PBS, pelleted, and resuspended in cold PBS.

For peripheral blood single cells, 100uL blood from mice were added into 5mL RBC lysis buffer (Tonbo Biosciences) and incubated at room temperature for 10 min. Cells were washed with cold PBS, pelleted, and process 10 min of RBC lysis once more. Cells were then washed again, pelleted, and resuspended in cold PBS.

### Flow cytometry

Single-cell suspensions were prepared from various mouse tissues, including the epidermis, and dermis, draining lymph nodes, spleens, blood, and bone marrow. For surface staining, cells were incubated in cold PBS for 30 min with the following flow antibodies: Zombie-violet (for live/death cells), anti-mouse CD45-BUV563, CD11b-BUV737, TCR γ/δ-APC, CD4-BUV395, CD8α-BV615, B220-AF700, F4/80-AF488, CD11c-BV711, NK1.1-PE-Cy5, Ly6G-Spark blue-550, CD3 ε-BV785, and CD16/32 (Fc blocker). For intracellular cytokine staining, cells were treated with a Cell Activation Cocktail (Biolegend) in RPMI-1640 medium containing 5% fetal bovine serum for 4 h at 37°C, followed by the surface staining using the following antibodies: CD45-BUV563, CD11b-BUV737, CD4-BUV395, CD8α-BV615, F4/80-AF488, CD11c-BV711, TCR γ/δ - PE-Cy5, CD3 ε-BV785, and CD16/32 (Fc blocker). Afterward, cells were washed with PBS, fixed, and permeabilized using the intracellular staining buffer set (Cat. #424401, Biolegend) according to the manufacturer’s instructions. Intracellular staining was performed using the following antibodies: anti-mouse TNFα-APC-Cy7, IFNγ-BV510, IL-17A-BV650, IL-1β-PE-Cy7, IL-4-BV605, and IL-10-Pacific Blue. For intracellular FABP5 staining, cells stained by surface markers underwent the intracellular staining process using goat anti-mouse FABP5 antibody and rabbit anti-mouse Karatin6A for 1.5 h at 4°C, followed by secondary staining of Donkey-anti-goat IgG-AF594 antibody and Donkey polyclonal anti-rabbit IgG-BV510 antibody. Cells were acquired by Cytek Aurora Flow Cytometer. Flow cytometric data analysis was performed using FlowJo v10 software.

### Analysis of bulk RNA sequencing datasets

RNA sequencing dataset (GSE161296) from our previous study [cite] was downloaded from the Gene Expression Omnibus (GEO) database. Relative expression levels of individual genes between mouse skins on LFD or cocoa butter HFD were extracted from the dataset. Differential gene expression (DEG) analysis was performed as previously described^17^. GSE151839 dataset^21^ consisting of extracted RNA from skin biopsies of 10 normal weight and 10 obese doners was analyzed using on-site GEO2R tool, an R based tool. Volcano plot and heatmap visualizations were performed using GraphPad Prism 10 software. Pathway analysis was conducted using g: Profiler [cite] to identify enriched pathways in the KEGG and Gene Ontology (GO: Biological Process) database.

### Analysis of single-cell RNA sequencing datasets

Single-cell RNA sequencing datasets of mouse skin (GSE132042)^22^ and human skin (GSE202352)^23^ were analyzed and visualized using the BxGenomics scRNA-Seq View (https://app.bxgenomics.com/bxg/app/). In both datasets, Fabp5^high^ and Fabp5^lo^ were identified based on Fabp5 gene expression level. Dot plots for gene expression levels UMAP visualizations, and Dual Gene analysis for gene co-expressions were perfoemed in BxGenomics platform. DEGs were selected based on an adjusted p-value <0.05 and log2 fold change ±0.26 and visualized as volcano plot using GraphPad Prism 10 software. Pathway analysis was conducted using Enrichr to identify enriched pathways in the Reactome Pathways database^24^.

### Cell culture

HaCaT cells were cultured in DMEM medium containing 10% FBS and antibiotics as previously described^9^. Primary epidermal single cells were obtained from Fabp5^f/f^ Krt6A-Cre mice and cultured in DMEM medium containing 10% FBS and antibiotics. To investigate the effect of FABP5 and MAPK signaling in saturated fatty acid (SFA)-induced CCL20 expression in keratinocytes, HaCat cells or primary epidermal cells were pretreated with SBFI-26 (10µM, FABP5 inhibitor), Adezmapimod (5µM, p-38 inihibitor), U0126 (10µM, ERK inhibitor) or Sp600125 (10µM, JNK inhibitor), for 1 hour, followed by 200µM stearic acid (SA) or BSA control for another 8 hours in the continued presence of inhibitors in DMEM medium containing 2% FBS. Cells were washed with cold PBS twice and lysed with RNA lysis buffer (Cat. #12183025, Invitrogen, Waltham, MA) for further RNA extraction.

### Quantitative real-time PCR

RNA was extracted from primary epidermal cells and HaCaT cells using PureLink RNeasy Mini Kit (Cat. #12183025, Invitrogen, Waltham, MA). The cDNA synthesis was performed with a QuantiTect Reverse Transcription Kit (Cat. #205314, Qiagen, Hilden, Germany) according to the manufacturer’s instruction. Real-time PCR was performed with Power SYBR Green PCR Master Mix (Cat. #4368708, Applied biosystems, Waltham, MA) using StepOnePlus qPCR Systems (Applied Biosystems) to analyze the expression of target genes encoding CCL20, FABP5, IL-23, IL-1β, IL36γ, IL-6, or IL-18 using primers from IDT (see primer sequences in Table S1). Housekeeping genes for murine and human were Hprt1 and GAPDH, respectively.

### PAGE and western blot

Skin tissue from Fabp5^f/f^ Krt6A-Cre mice were lysis with RIPA Buffer (Cat. #9806S, Cell Signaling Technology). Proteins were extracted from the lysate by centrifugation at maximum speed, followed by supernatant collection. Protein concentrations were measured in each sample using BCA assay (Cat. #23225, Thermo Scientific). The samples were then mixed with SDS loading buffer containing 2-mercaptoethanol, boiled at 90°C for 5 min, and loaded to an SDS-PAGE gel to be separated. After SDS-PAGE, proteins were transferred to polyvinylidene difluoride membranes. Target proteins were probed with specific antibodies, including anti-mouse Phospho-p38 (Cat. #9216, Cell Signaling Technology), p38 (Cat. #9228, Cell Signaling Technology), and FABP5 (Cat#AF1476, R&D Systems) with β-actin (Cat. # MA1-140-A488, Thermo Scientific) as the loading control. A Bio-Rad imaging system was used to visualize protein levels.

### Quantification and statistical analysis

All data were presented as the mean ± SD. All experiments were performed by at least three independent replicates. For both *in vitro* and *in vivo* experiments, two-tailed, unpaired Student’s t-test, two-tailed unpaired multiple t test, one-way ANOVA followed by Bonferroni’s multiple comparisons test, and two-way ANOVA with Tukey’s multiple comparison test were used.

## Results

### FABP5 deficiency in keratinocytes attenuates saturated high-fat diet (HFD)-exacerbated psoriatic inflammation in mice

To determine the role of FABP5 in keratinocyte on saturated HFD-induced obesity, we generated keratinocyte FABP5 conditional knockout mice (Fabp5^f/f^ Krt6a-Cre) as previously described ^9^, and validated the efficiency of FABP5 deficiency in skin keratinocytes by both Western blotting and flow cytometry staining (Figure 1a and b). To determine whether keratinocyte-specific FABP5 deficiency affects adiposity, we fed the Fabp5^f/f^ Krt6a-Cre mice with saturated HFD (cocoa butter, 45% fat) or low-fat diet (LFD, 5% fat) for 5 months as previously described^11^. Compared with LFD control, saturated HFD induced comparable increases in body weight (Figure 1c) and fat mass (Figure 1d), without altering lean mass (Figure 1e), in both Fabp5^f/f^ Krt6a-Cre^-^ and Fabp5^f/f^ Krt6a-Cre^+^ mice. These results indicate that keratinocyte-specific FABP5 deficiency does not affect mouse obese status by saturated HFD feeding.

**Figure 1.**
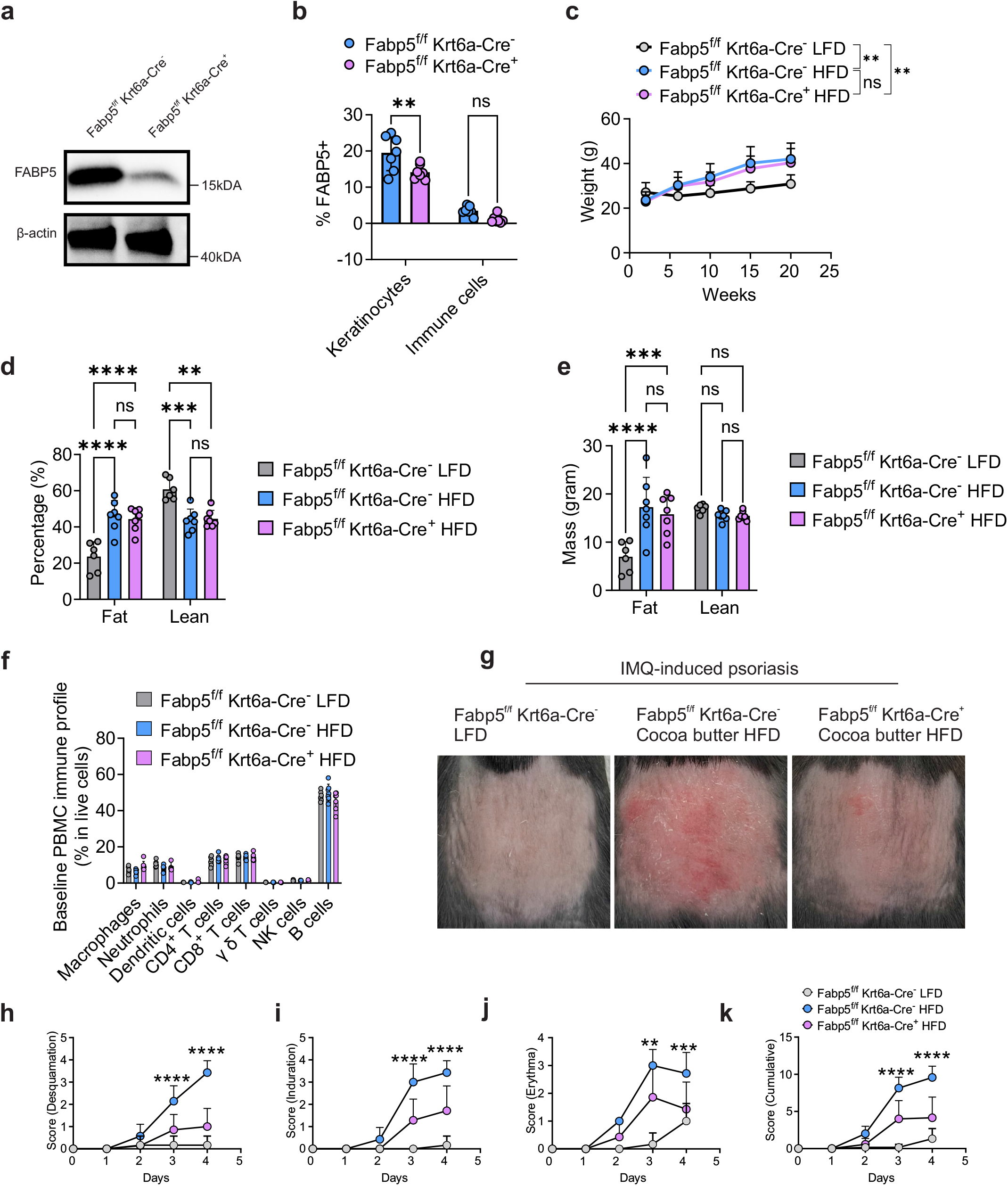
Keratinocyte-specific FABP5 deficiency attenuates saturated high-fat diet (HFD)-exacerbated psoriatic inflammation in mice. (a) Western blotting of FABP5 expression in fat-removed skin tissue of Fabp5^f/f^ Krt6a-Cre^-^ and Fabp5^f/f^ Krt6a-Cre^+^ mice. (b) Flow cytometric analysis of intracellular FABP5 in live epidermal keratinocytes (Zombie^-^ CD45^-^ Krt6A^+^) and immune cells (Zombie^-^ CD45^+^). (c) Body weight curve of Fabp5^f/f^ Krt6a-Cre mice fed LFD or saturated HFD. (d-e) NMR scanning for percentages (d) and mass (e) of body fat and lean content of Fabp5^f/f^ Krt6a-Cre mice fed saturated HFD or LFD for 5 months. (f) Flow cytometric analysis of peripheral blood immunophenotypes of Fabp5^f/f^ Krt6a-Cre fed saturated HFD or LFD for 5 months. (g) Representative images of skin from Fabp5^f/f^ Krt6a-Cre mice fed saturated HFD or the LFD for 5 months, followed by 4 days of IMQ treatment (4 mg/mouse). (h-k) Evaluation of erythema (h), induration (i), and desquamation (j) from day 0 of IMQ treatment in mice on customized HFDs versus the LFD, and the cumulative score (k) summing from (h-j). Data are shown as mean ± SD, statistical significance was determined using two-way ANOVA with Tukey’s multiple comparison test. **, P < 0.01; ***, P < 0.001; ****, P < 0.0001; ns, nonsignificant.

To assess whether keratinocyte-specific FABP5 deficiency influences obesity-associated systemic immune regulation, we next analyzed baseline peripheral blood immune profiles in Fabp5^f/f^ Krt6a-Cre mice fed saturated HFD or LFD control (Figure S1a). Under the saturated HFD, Fabp5^f/f^ Krt6a-Cre^+^ mice exhibited minimal changes in peripheral immune populations compared with Fabp5^f/f^ Krt6a-Cre^-^ mice (Figure 1f). Functional analyses similarly showed that saturated HFD feeding had little effect on the production of common inflammatory cytokines by major immune cell populations, including monocytes, CD4, CD8+ and *γδ* T cells, with the exception of modestly increased TNFα production by CD4^+^ and CD8^+^ T cells. Notably, no significant differences in cytokine production were observed between Fabp5^f/f^ Krt6a-Cre^-^ and Fabp5^f/f^ Krt6a-Cre^+^ mice fed the saturated HFD (Figure S1b-S1e). Together, these findings suggest keratinocyte-specific FABP5 deficiency has minimal impact on the systemic immune profiles under saturated HFD feeding.

To determine whether keratinocyte-specific FABP5 contributes to saturated HFD-related psoriasis, we applied imiquimod (IMQ) on the shaved dorsal skin of the Fabp5^f/f^ Krt6a-Cre mice fed the saturated HFD or LFD. Consistent with our previous findings^11^, saturated HFD accelerated the onset of IMQ-induced psoriatic skin inflammation in Fabp5^f/f^ Krt6a-Cre^-^ (WT) mice. Notably, the saturated HFD-exacerbated psoriatic inflammation was significantly ameliorated in Fabp5^f/f^ Krt6a-Cre^+^ mice (Figure 1g). Further analysis of psoriatic severity, evaluated by desquamation, induration, erythema, and the cumulative scores, showed that saturated HFD-increased psoriatic symptoms in Fabp5^f/f^ Krt6a-Cre^-^ mice were significantly attenuated in Fabp5^f/f^ Krt6a-Cre^+^ mice (Figure 1h-k). Collectively, these findings demonstrate that FABP5 expression in keratinocytes contributes to saturated HFD-exacerbated psoriatic inflammation.

### Keratinocyte-specific FABP5 deficiency reduced proinflammatory γ δ T-cell accumulation in saturated HFD-induced psoriatic inflammation

To investigate how keratinocyte-specific FABP5 deficiency attenuated saturated HFD-exacerbated psoriatic inflammation, we performed a comprehensive flow cytometric analysis of immune cell profiles across multiple tissues in saturated HFD-induced psoriatic models (Figure 2a, Figure S1a, S2a-S2d). Consistent with our previous findings^11^, saturated HFD increased the proportions of proinflammatory TCR γ δ ^dim^ cells in both epidermis and dermis of psoriasis compared to LFD in Fabp5^f/f^ Krt6aCre^-^ mice. Notably, these saturated HFD-induced increases were significantly attenuated in Fabp5^f/f^ Krt6aCre^+^ mice (Figure 2b–2d), suggesting that keratinocyte-specific FABP5 deficiency suppresses saturated-HFD induced accumulation of proinflammatory TCR γ δ ^dim^ cell in the psoriatic skin. In contrast, the proportions of other major immune subsets in epidermis, dermis, or draining lymph nodes (dLNs) remained comparable between Fabp5^f/f^ Krt6aCre^-^ and Fabp5^f/f^ Krt6aCre^+^ mice (Figure 2e–2g). Similarly, no differences in these immune subsets were observed in systemic immune compartments, including peripheral blood and spleen, between two groups fed saturated HFD (Figure S3a). These findings indicate that keratinocyte-specific FABP5 selectively promotes the local accumulation of proinflammatory TCR γ δ ^dim^ cells in psoriatic skin, rather than broadly altering local or systemic immune cell composition in response to the saturated HFD feeding. Collectively, these identify keratinocyte FABP5 as an important mediator linking saturated HFD to psoriatic inflammation by promoting accumulation of proinflammatory TCR γ δ ^dim^ cells in the skin.

**Figure 2.**
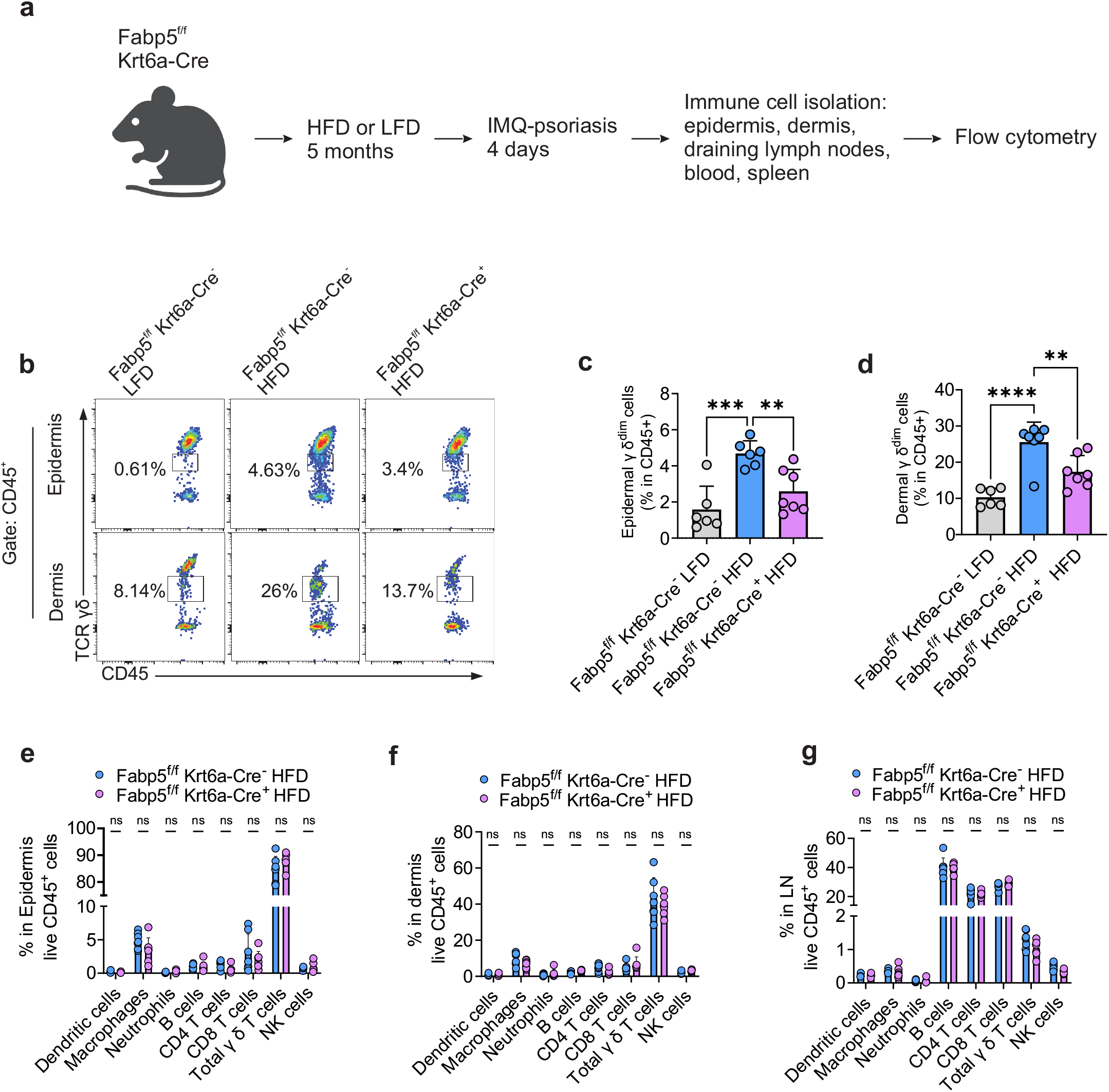
Keratinocyte-specific FABP5 deficiency reduced proinflammatory γ δ T-cell accumulation in psoriatic skin under saturated HFD. (a) Schematic workflow of comprehensive multi-organ immunophenotyping from Fabp5^f/f^ Krt6a-Cre mice fed saturated HFD or the LFD for 5 months, followed by 4 days of IMQ treatment. (b) Representative flow cytometric gating for TCR γ δ ^dim^ cells under gate CD45^+^Zombie^−^CD11b^-^ in the psoriatic epidermis and dermis from mice Fabp5^f/f^ Krt6a-Cre mice fed saturated HFD or the LFD for 5 months. (c-d) Statistical analysis of ratios of TCR γ δ ^dim^ cells in CD45^+^Zombie^−^ live immune cells in the psoriatic epidermis (c) and dermis (d) from mice Fabp5^f/f^ Krt6a-Cre mice fed saturated HFD or the LFD for 5 months by flow cytometry. (e-g) Statistical analysis of Immunophenotype of epidermis (e), dermis (f), and draining lymph nodes (dLN) (g) from mice Fabp5^f/f^ Krt6a-Cre mice fed saturated HFD or the LFD for 5 months by flow cytometry. Data are shown as mean ± SD, statistical significance was determined using one-way ANOVA with Bonferroni’s multiple comparison test for (c-d) and two-way ANOVA with Tukey’s multiple comparison test for (e-g). **, P < 0.01; ***, P < 0.001; ****, P < 0.0001; ns, nonsignificant.

### Keratinocyte-specific FABP5 deficiency reduces IL-17A production by γ δ T cells in saturated HFD-induced psoriasis

To further investigate the role of keratinocyte FABP5 in regulating immune cell function in saturated HFD-exacerbated psoriasis, we analyzed cytokine production by immune cells from psoriatic skin and dLNs. Intracellular cytokine staining followed by flow cytometric analysis revealed that saturated HFD increased IL-17A production by total γ δ T cells in epidermis, dermis, and dLNs compared with LFD in Fabp5^f/f^ Krt6aCre^-^ mice (Figure 3a). However, these HFD-induced increases in IL-17A production in epidermis, dermis and dLNs were significantly attenuated in Fabp5^f/f^ Krt6aCre^+^ mice (Figure 3b–3d). This pattern was also observed in TCR γ δ dim cells in both dermis and epidermis (Figure S3b) indicating that keratinocyte FABP5 contributes to the enhanced IL-17A responses of γ δ T cells under the saturated HFD. In contrast, keratinocyte-specific FABP5 deficiency had minimal effects on the production of other cytokines by γ δ T cells, including IFNγ, IL-4, TNFα, and IL-10 (Figure 3e–3g). Similarly, FABP5 deficiency in keratinocytes exhibited no effect on cytokine production in CD4^+^ T cells, CD8^+^ T cells, and macrophages in the epidermis, dermis, and dLNs in saturated HFD-related psoriasis (Figure S3c-S3e). These results suggest that the effect of keratinocyte FABP5 on immune cell function in HFD-induced psoriatic inflammation is selective for the IL-17A response of γ δ T cells rather than reflecting broad alterations in inflammatory cytokine production. Taken together, these findings demonstrate that keratinocyte FABP5 selectively promotes saturated HFD-enhanced IL-17A production by γ δ T cells, supporting a keratinocyte–γ δ T-cell axis through which dietary saturated fat exacerbates psoriatic inflammation.

**Figure 3.**
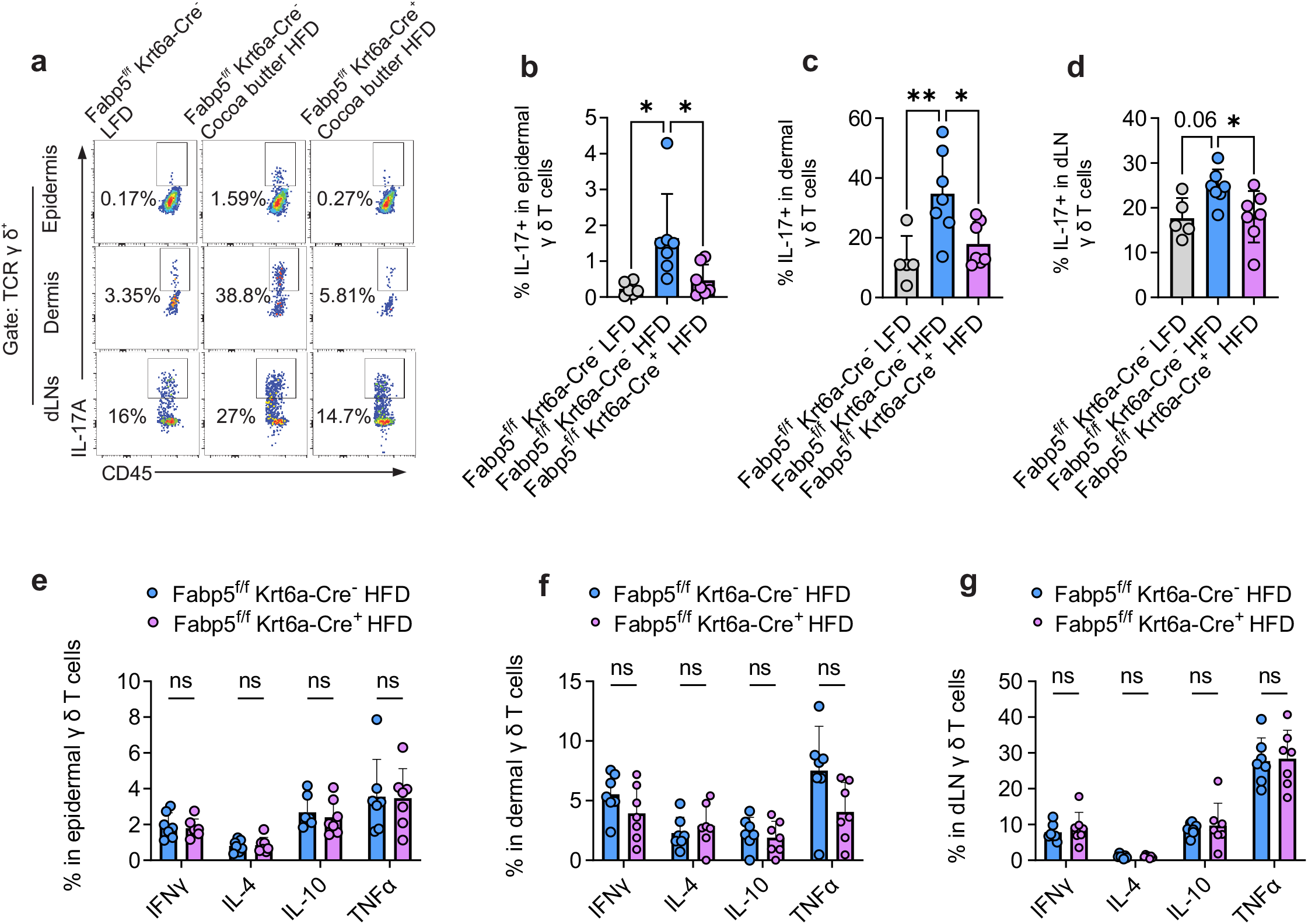
Keratinocyte-specific FABP5 deficiency reduces IL-17A production by γ δ T cells in psoriatic skin under saturated HFD. (a) Representative flow cytometric gating for IL-17A^+^ in TCR γ δ^+^ cells in epidermis, dermis, and dLNs from Fabp5^f/f^ Krt6a-Cre mice fed saturated HFD or the LFD for 5 months, followed by 4 days of IMQ treatment. (b-d) Statistical analysis of IL-17A^+^ ratio in γ δ T cells in epidermis (b), dermis (c), and dLNs (d) by flow cytometry. (e-g) Statistical analysis of ratios of IFNγ^+^, IL-4^+^, IL-10^+^, and TNFα^+^ in γ δ T cells in epidermis (e), dermis (f), and dLNs (g) by flow cytometry. Data are shown as mean ± SD, statistical significance was determined using one-way ANOVA with Bonferroni’s multiple comparison test for (b-d) and two-way ANOVA with Tukey’s multiple comparison test for (e-g). *, P< 0.05, **, P < 0.01; ***, ns, nonsignificant.

### Saturated HFD is associated with enhanced proinflammatory MAPK signaling in skin

To determine the molecular mechanisms by which saturated HFD promotes proinflammatory signaling in skin keratinocytes for γ δ T-cell activation, we first analyzed bulk RNA-sequencing dataset from mice fed saturated HFD or LFD control (GSE161296)^25^. As illustrated by the volcano plot and heatmap, saturated HFD upregulated genes related to fatty acid metabolism (e.g., *Cd36, Pparg, Fabp4*) and multiple genes related to MAPK pathways (e.g., *Fgf1, Fos, Mef2c, Fgf6, Gadd45g, Pdgfb, Mapk12, and Ereg*) compared with LFD control (Figure 4a and 4b). Notably, *Rorc*, which encodes the transcriptional factor RORγt and is critical for IL-17-producing γ δ T cells in psoriatic inflammation^26^, was also upregulated in the skin of saturated HFD-fed mice, consistent with an enhanced cutaneous IL-17-associated inflammatory environment (Figure 4a, 4b). Kyoto Encyclopedia of Genes and Genomes (KEGG) pathway enrichment analysis further identified enrichment of pathways related to fatty acid metabolism and MAPK signaling pathways (Figure 4c). These data suggest that saturated HFD is associated with enhancement of lipid metabolic and proinflammatory MAPK programs in the mouse skin.

**Figure 4.**
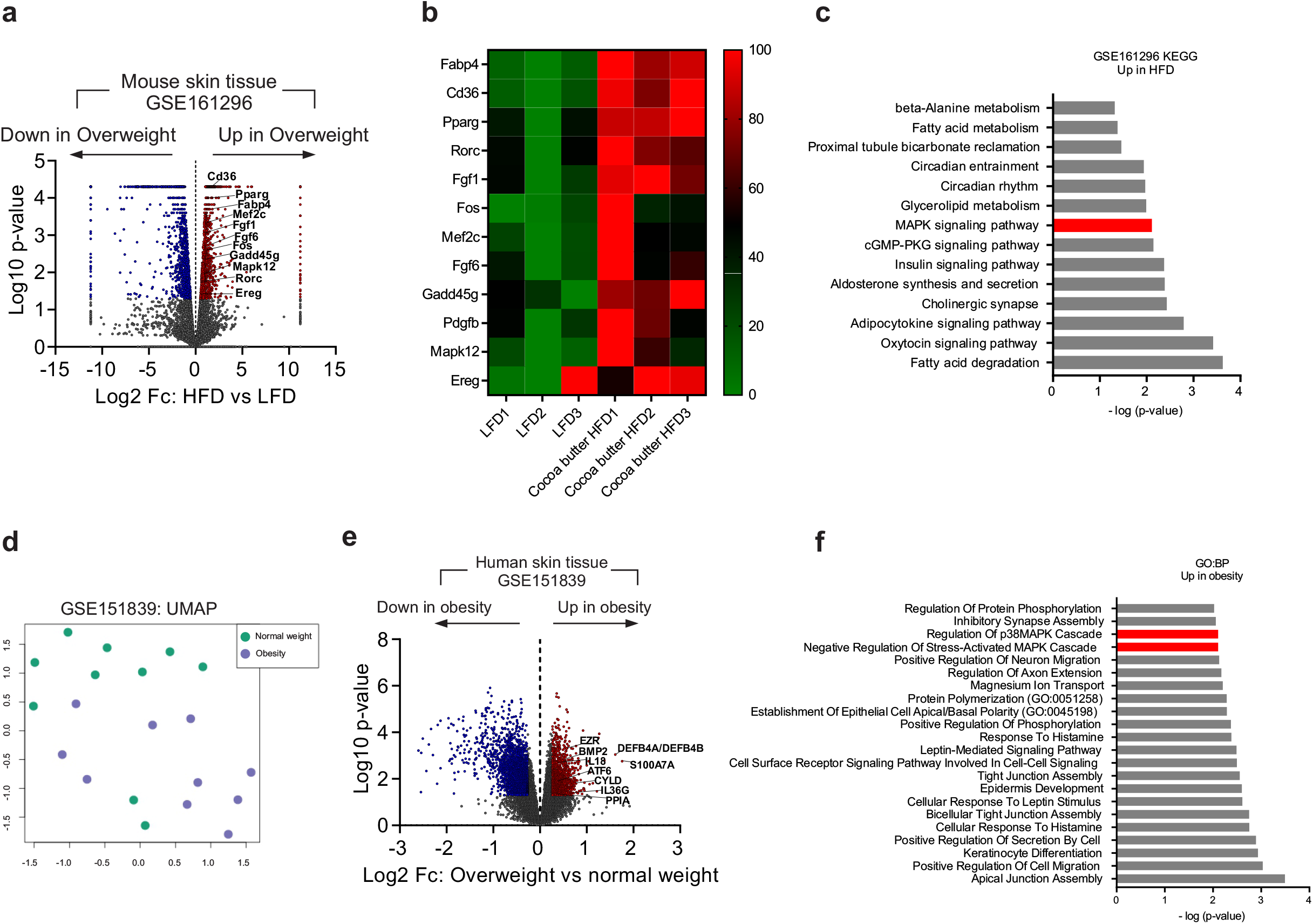
Saturated HFD is associated with enhanced proinflammatory MAPK signaling in skin. (a) Volcano plot showing differentially expressed genes (DEGs) identified by bulk RNA sequencing of whole-skin samples from mice fed a saturated HFD or LFD, using a p-value cutoff of 0.05 and a Log_2_Fc cutoff of ±0.26. Red dots are significantly upregulated genes in saturated HFD group, blue dots are significantly downregulated genes in saturated HFD group, and gray dots are genes with no significant difference. (b) Heatmap of significantly upregulated genes in whole-skin of mice fed a saturated HFD compared to LFD. (c) Pathway analysis using the KEGG database of DEGs upregulated in the saturated HFD group. (d) Uniform Manifold Approximation and Projection (UMAP) analysis of gene expression between skin biopsies from normal-weight (n=10) and overweight (n=10) donors in bulk RNA-sequencing. (e) Volcano plots showing DEGs identified by bulk RNA sequencing of skin biopsies from normal-weight and obese donors, using a p-value cutoff of 0.05 and a Log2Fc cutoff of ±0.26. Red dots are significantly upregulated genes in obesity group, blue dots are significantly downregulated genes in obesity group, and gray dots are genes with no significant difference. (f) Analysis of gene oncology biological process (GO:BP) from DEGs upregulated in obesity group.

To determine whether similar molecular alterations are present in human skin in the context of obesity, we next analyzed human skin bulk RNA-sequencing dataset (GSE151839)^21^. Uniform Manifold Approximation and Projection (UMAP) analysis revealed distinct clustering of skin samples from normal weight (BMI 18.5-26.9) and obesity (BMI 35-50), suggesting substantial differences in cutaneous gene-expression profiles between two groups (Figure 4d). Consistent with mouse dataset, multiple MAPK-related genes, including EZR, BMP2, CYLD, and PPIA, were upregulated in the skin of obese individuals, as illustrated by volcano plot (Figure 4e) and pathway enrichment using Gene Ontology (GO) analysis (Figure 4f). In addition, genes related to psoriatic inflammation (e.g., *IL18, IL36G*), antibacterial peptides (e.g., *DEFB4A/DEFB4B, S100A7A*), as well as ER-stress-related marker, *ATF6*, were upregulated in the skin from obese patients (Figure 4e). These transcriptional changes are consistent with an enhanced inflammatory and cellular stress environment in the skin associated with obesity. Taken together, analyses of mouse and human skin transcriptomic datasets reveal a conserved association between saturated HFD feeding/adiposity and enhanced cutaneous MAPK-related inflammatory programs. These findings support MAPK signaling as a potential pathway linking saturated HFD-associated metabolic alterations to a proinflammatory skin environment.

### FABP5 mediates saturated fat-induced proinflammatory MAPK signaling in keratinocytes

Given the critical role of keratinocyte-specific FABP5 in meditating saturated HFD-exacerbated psoriatic inflammation (Figure 1–3) and the close link between obesity and cutaneous MAPK programs (Figure 4), we hypothesized that FABP5 promotes saturated fat-induced inflammatory MAPK signaling in keratinocytes. To address this, we first analyzed mouse skin single-cell RNA sequencing dataset (GSE132042)^22^. Clustering analysis based on established cell-type marker genes identified 7 distinct cell populations in the mouse skin (Figure 5a, S4a). Within the keratinocyte compartment, we identified two distinct subsets characterized by relatively high or low *Fabp5* expression (Fabp5^high^ and Fabp5^low^, respectively) (Figure 5b). Consistent with the lipid chaperon function of FABP5, Fabp5^high^ keratinocytes exhibited increase expression of genes related to mitochondrial fatty-acid β-oxidation (e.g., *Acaa2, Acadm, Hadh*, etc.) (Figure S4b), suggesting an enhanced capacity for fatty acid handling and mitochondrial lipid metabolism.

**Figure 5.**
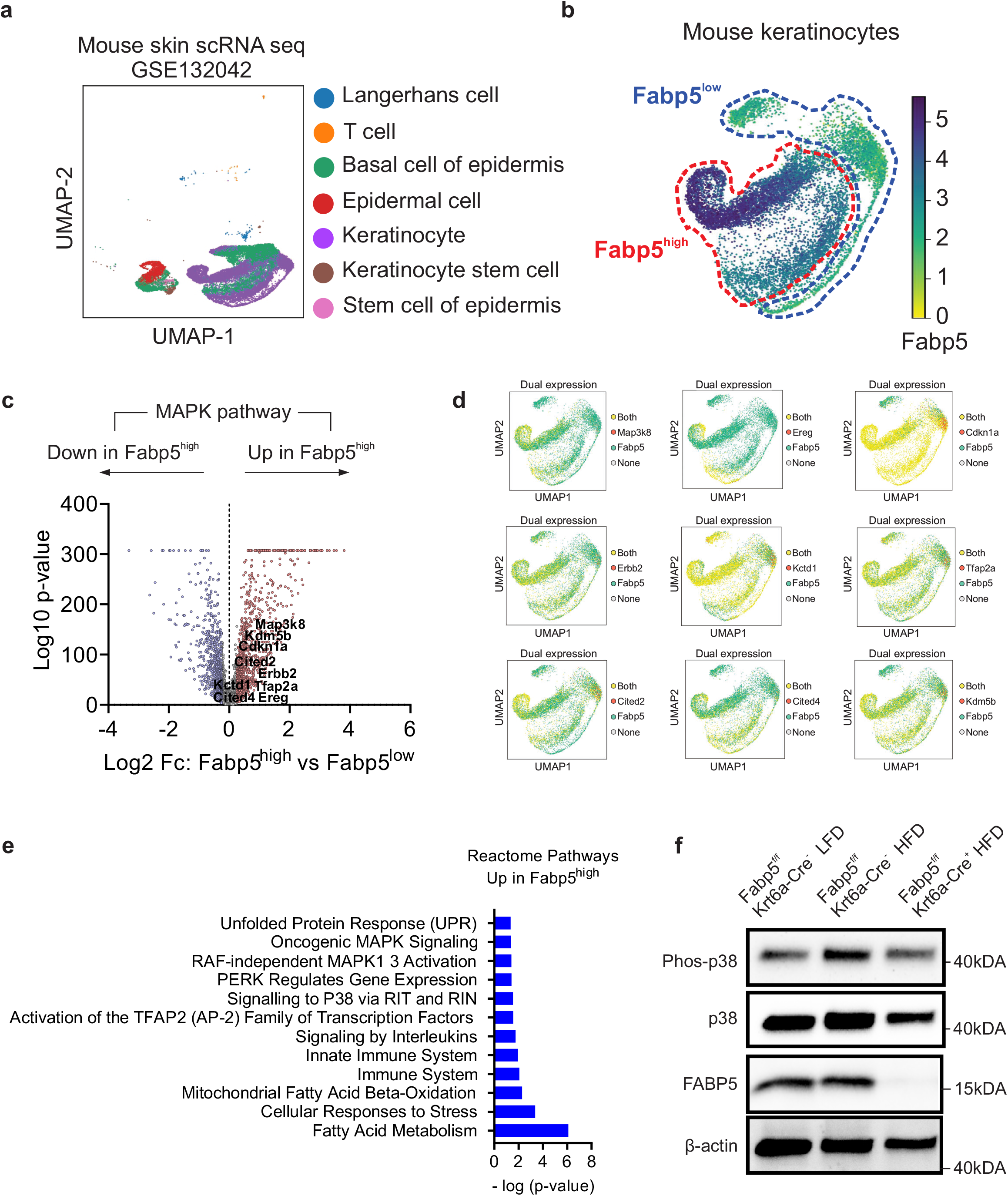
FABP5 mediates saturated fat-induced proinflammatory MAPK signaling in keratinocytes. (a) Single-cell RNA sequencing (ScRNA-seq) clustering analysis of mouse skin tissue. (b) Identification of Fabp5^high^ and Fabp5^low^ mouse keratinocytes based on Fabp5 gene expression level in ScRNA-seq. (c) Volcano plot of DEGs involved in MAPK pathways by comparing Fabp5^high^ and Fabp5^low^ keratinocytes using an adjusted p-value cutoff of 0.05 and a Log2FC cutoff of ±0.26. Red dots are significantly upregulated genes in Fabp5^high^, blue dots are significantly downregulated genes in Fabp5^high^, and gray dots are genes with no significant difference. (d) Dual-gene expression analysis of Fabp5 and MAPK-related genes in keratinocytes visualized by UMAP showing cells expressing Fabp5 alone, MAPK-related genes alone, both Fabp5 and MAPK-related genes, or neither gene (none). (e) Pathway enrichment analysis of significantly upregulated genes in Fabp5^high^ keratinocytes using Reactome Pathways database. (f) Western blot analysis of total p38, phosphorylated p38, phosphorylated JNK, and FABP5 protein levels in the lysate of fat-removed skin from Fabp5^f/f^ Krt6a-Cre mice fed saturated HFD or LFD.

Beyond metabolic phenotype, Fabp5^high^ keratinocytes exhibited increased expression of genes related to ER stress (e.g., *Atf3, Atf4, Atf5*, ect.) (Figure S4c), as well as genes involved in psoriasis-associated inflammatory responses (e.g., *Ccl20, Pycard, Il17rc, Nfkbia, Myd88, Il20rb, Il22ra1, S100a7a*, etc.) (Figure S4d). Notably, multiple genes associated with MAPK signaling (e.g., *Map3k8, Ereg, Cdkn1a, Erbb2, Cited2, Cited4, Tfap2a*, etc.) were elevated in Fabp5^high^ keratinocytes (Figure 5c). Dual expression analysis further demonstrated co-expressions of *Fabp5* with multiple MAPK pathway-related genes, including *Map3k8, Ereg, Cdkn1a, Erbb2, Kctd1, Tfap2a, Cited2, Cited4, and Kdm5b* (Figure 5d). Reactome Pathway analysis showed enrichment of pathways associated with lipid metabolism, ER stress, immune activation, and MAPK signaling in Fabp5^high^ keratinocytes (Figure 5e). To determine whether a similar phenotype exists in human skin, we analyzed human skin scRNA-sequencing dataset (GSE202352)^23^ and compared FABP5^high^ and FABP5^low^ keratinocytes (Figure S4e-S4g). Consistent with the mouse dataset, human FABP5^high^ keratinocytes exhibited enhanced expression of genes associated with MAPK signaling, mitochondrial fatty acid oxidation, and immune activation (Figure S4h and S4i). Collectively, these analyses identify FABP5^high^ keratinocytes as a metabolically active, proinflammatory population characterized by enhanced lipid metabolic and MAPK-associated transcriptional programs.

To further determine whether keratinocyte FABP5 contributes to saturated HFD-induced MAPK activation *in vivo*, we isolated proteins from skin of LFD- or HFD-fed Fabp5^f/f^ Krt6a-Cre mice and assessed MAPK signaling by Western blotting. Saturated HFD increased phosphorylated p38 (p-p38) and total p38 expression in the skin of Fabp5^f/f^ Krt6a-Cre^-^, whereas these increases were absent or attenuated inFabp5^f/f^ Krt6a-Cre^+^ mice (Figure 5f). Together, these findings support a critical role for keratinocyte FABP5 in mediating saturated HFD-induced MAPK pathway activation in the skin, providing a molecular link between FABP5-dependent lipid sensing and proinflammatory MAPK signaling in keratinocytes.

### Keratinocyte FABP5 links saturated fat-induced γ δ T-cell IL-17 responses through the MAPK/CCL20 signaling

C-C motif chemokine ligand 20 (CCL20) is a well-established keratinocyte-derived chemokine regulated by MAPK pathways that promotes the activation and recruitment of IL-17-producing γ δ T cells in the skin^27, 28^. Given the essential role of keratinocyte-specific FABP5 in meditating saturated HFD-enhanced IL-17A in skin γ δ T cells (Figure 3), we hypothesize that keratinocyte FABP5 links saturated fat exposure to γ δ T-cell IL-17 responses through the MAPK/CCL20 signaling. To test this, we first analyzed *Ccl20* expression in mouse skin scRNA-sequencing dataset and found that *Ccl20* was enriched in Fabp5^high^ keratinocytes and showed strong co-expression with *Fabp5* (Figure 6a–6c), suggesting a potential association between FABP5 and CCL20 signaling in keratinocytes.

**Figure 6.**
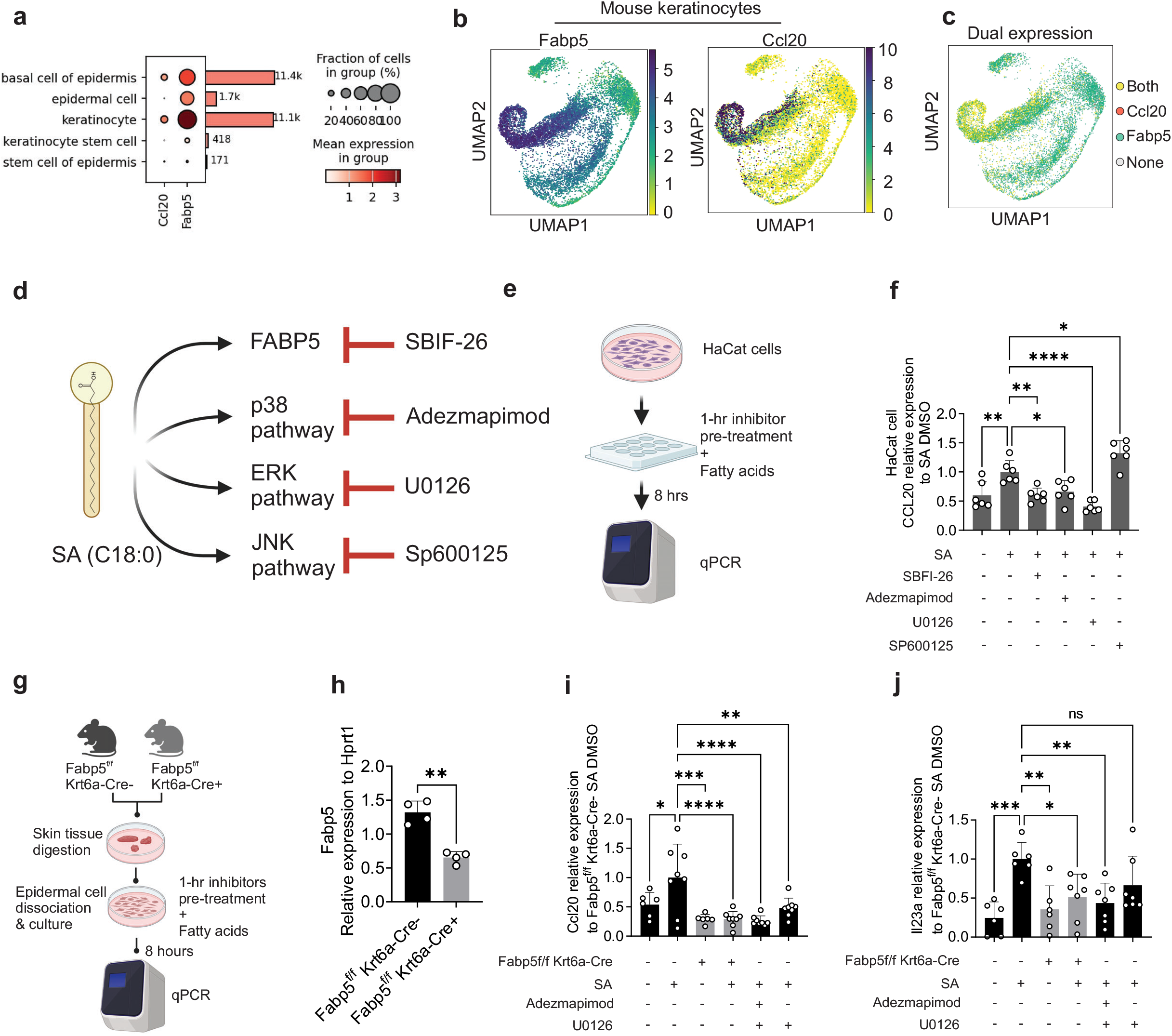
Saturated fat promotes CCL20 in keratinocytes via FABP5/MAPK axis. (a) Dot plot of Fabp5 and Ccl20 gene expression across non-immune cell subsets in the mouse skin by scRNA-seq. (b) UMAP analysis of Fabp5 and Ccl20 gene expression in mouse keratinocytes by scRNA-seq. (c) Dual-gene expression analysis of Fabp5 and Ccl20 in keratinocytes visualized by UMAP showing cells expressing Fabp5 alone, Ccl20 alone, both Fabp5 and Ccl20, or neither gene (none). (d) Schematic of determination of pathways involved in SA-induced CCL20 expression. (e) Schematic of qPCR assay on HaCat cells pretreated different inhibitors for 1-hours followed by 8 hours of fatty-acid treatment in the continued presence of inhibitors. (f) Expression levels of CCL20 by qPCR in HaCat pretreated different inhibitors for 1-hours followed by 8 hours of fatty-acid treatment in the continued presence of inhibitors. (g) Schematic of primary epidermal single cell isolation from Fabp5^f/f^ Krt6a-Cre mice, followed by 1-hour inhibitor pretreatment and 8-hour fatty acid treatment in the continued presence of inhibitors, with gene expression assessed by qPCR at the endpoint. (h) Expression levels of Fabp5 by qPCR in the primary epidermal single cell isolated from Fabp5^f/f^ Krt6a-Cre mice. (i, j) Expression levels of Ccl20 (i) and Il23a (j) by qPCR in primary keratinocytes isolated from Fabp5^f/f^ Krt6a-Cre mice followed by 1-hour inhibitor pretreatment and 8-hour fatty acid treatment in the continued presence of inhibitors.

To determine whether the FABP5/MAPK axis directly regulates CCL20 expression in response to saturated fat, we treated HaCat cells, a human keratinocyte cell line, *in vitro* with BSA control or stearic acids (SA) in the presence or absence of inhibitors targeting FABP5 (SBIF-26), p38 (Adezmapimod), ERK (U0126), or JNK (Sp600125), followed by qPCR analysis (Figure 6d and 6e). SA treatment significantly increases *CCL20* mRNA expression compared with BSA control (Figure 6f), demonstrating that saturated fatty acid exposure induces CCL20 expression in keratinocytes. Importantly, pharmacological inhibition of FABP5, p38, or ERK significantly attenuated SA-induced *CCL20* expression, whereas JNK inhibition with SP600125 showed no inhibition effect (Figure 6f). These results indicate that SA-induced CCL20 expression requires FABP5 and is mediated predominantly through the MAPK signaling. We next examined whether this regulatory mechanism extended to other inflammatory mediators. Although SA induced expressions of proinflammatory cytokines, such as *IL1B*, *IL36G*, and *IL18*, inhibition of FABP5, p38, ERK, or JNK did not significantly attenuate these responses (Figure S5a-S5c), suggesting that their induction is independent of the FABP5/MAPK pathway. Together, these results identify the FABP5/MAPK axis as a selective mechanism mediating saturated fat-induced CCL20 expression in human keratinocytes.

To validate this mechanism in primary mouse keratinocytes, we isolated keratinocytes from Fabp5^f/f^ Krt6a-Cre mice and treated them *in vitro* with BSA control or SA in the presence or absence of MAPK inhibitors, followed by qPCR analysis (Figure 6g and 6h). Consistent with our findings in HaCat cells, SA significantly induced *Ccl20* expression in Fabp5^f/f^ Krt6a-Cre^-^ keratinocytes, whereas this induction was markedly attenuated in Fabp5^f/f^ Krt6a-Cre^+^ keratinocytes (Figure 6i). In contrast, SA did not significantly induce *Il36g* or *Il6* expression in primary mouse keratinocytes (Figure S5d and S5e). Furthermore, inhibition of either p38 with Adezmapimod or ERK with U0126 attenuated SA-induced *Ccl20* expression in Fabp5^f/f^ Krt6a-Cre^-^ keratinocytes (Figure 6i), further supporting a FABP5-dependent MAPK mechanism underlying saturated fat-induced CCL20 expression. Notably, SA induced *Il23a* expression in Fabp5^f/f^ Krt6a-Cre^-^ keratinocytes, and this response was attenuated by keratinocyte-specific FABP5 deficiency and p38 inhibition (Figure 6j), suggesting that FABP5/MAPK signaling may additionally enhance IL-17-associated inflammatory responses through IL-23 in mouse keratinocytes.

Collectively, these findings establish a mechanistic pathway whereby saturated fat is sensed through keratinocyte FABP5 to activate p38/ERK MAPK signaling and induce CCL20 expression. This FABP5/MAPK/CCL20 axis provides a mechanistic link between dietary saturated fat and enhanced recruitment and IL-17A response of proinflammatory γ δ T cells in psoriatic skin.

## Discussion

First described in antiquity, psoriasis is now increasingly recognized as a systemic inflammatory disease closely associated with metabolic dysfunction, particularly in the context of diet-induced obesity^29–31^. Among different types of dietary fats, SFAs have been selectively associated with exacerbated psoriatic inflammation^7, 10^. However, the cellular and molecular mechanisms by which increased SFA availability and metabolism in obesity reshape cutaneous immune responses remain to be determined.

In this study, we identified a previously unappreciated FABP5/MAPK/CCL20 axis that links saturated fats to psoriatic inflammation. In mouse models, keratinocyte-specific FABP5 deficiency attenuated saturated HFD-exacerbated psoriasis without affecting HFD-induced adiposity. Skin immune phenotyping and functional analyses demonstrated FABP5 deletion in keratinocytes reduced accumulation of proinflammatory TCR γ δ ^dim^ cells and decreased IL-17A production by these cells in saturated HFD-related psoriasis. Transcriptomic analyses of both mice and humans suggested that keratinocyte FABP5 expression is strongly associated with enhanced MAPK signaling under saturated HFD/obesity conditions. Importantly, cellular studies further demonstrated that SFAs induce upregulation of CCL20, a well-known chemokine that attracts IL-17-producing γ δ T cells through the FABP5-dependent MAPK pathway in keratinocytes.

As the predominant FABP family member expressed in the skin, FABP5 is highly abundant in keratinocytes and plays an important role in keratinocyte differentiation and functions^32, 33^. Our previous study demonstrated a proinflammatory function of FABP5 by forming a complex with VCP to promote NF-κB signaling in keratinocytes in non-obese mice^9^. The current study further extended these observations by showing FABP5 as a metabolic sensor linking increased SFA availability to inflammatory signaling in keratinocytes. We observed enhanced MAPK pathway activity in the skin under saturated HFD/obesity and a strong association between FABP5 expression and MAPK-related transcriptional programs in keratinocytes. Moreover, keratinocyte-specific FABP5 deficiency attenuated saturated fat-induced MAPK activation in mouse skin. Given that MAPK pathways are closely associated with obesity and can be activated by multiple forms of metabolic stress, including oxidative and endoplasmic reticulum (ER) stress^34, 35^, our findings support the role of keratinocyte FABP5 as a metabolic hub that translates lipid cues into inflammatory signals.

Importantly, these mechanisms provide a knowledge advance into how lipids reshape keratinocyte-immune cells crosstalk in the pathogenesis of psoriasis^36, 37^. Excessive lipids can profoundly influence tissue-resident cellular networks and have been closely linked to enhanced IL-17 responses in tissue inflammation^38, 39^. Our prior study has demonstrated that FABP5 in skin macrophages mediates SFA-induced IL-1β signaling, therefore promoting downstream IL-17 responses in saturated HFD-related psoriasis^7^. The current findings further identified a keratinocyte-dependent mechanism in which FABP5 senses SFAs and enhances skin γ δ T cell IL-17 responses in psoriatic inflammation. Importantly, we observed a strong correlation between FABP5 and CCL20 expression in keratinocytes by single-cell RNA sequencing analysis. Pharmacological FABP5 inhibition and FABP5 deficiency reduced SFA-induced CCL20 expression in keratinocytes, suggesting the FABP5/CCL20 axis in the recruitment and activation of IL-17-producing γ δ T cells in the skin^40^. Collectively, these findings reveal a cell-specific but convergent functions of FABP5 in keratinocytes and macrophages, establishing FABP5 as a key molecular interface between saturated fat handling and IL-17-driven inflammation in obesity-related psoriasis.

Notably, the conventional framework of “metaflammation” describes chronic, low-grade inflammation arising in response to nutrient excess and metabolic stress^41^. Classical models of obesity-associated metaflammation have focused primarily on adipocytes and immune cells, particularly macrophages, as key sensors and effectors of nutrient excess that contribute to systemic inflammatory and metabolic dysfunction^42^. Interestingly, our data showed that keratinocyte FABP5 conditional knockout did not alter adiposity or substantially affect systemic immune profiles under saturated HFD, but keratinocyte-specific FABP5 deficiency markedly protected mice against saturated HFD-exacerbated local IL-17 response and psoriatic inflammation. These findings suggest that metabolic inflammation can be initiated and regulated at the tissue level independently of detectable changes in systemic metabolic or immune status. Thus, beyond the classical adipocyte–immune cell paradigm of systemic metaflammation, our study suggests keratinocytes as tissue-resident metabolic sensors capable of translating excess lipid availability into a localized inflammatory program. This concept of tissue-localized metaflammation may provide a framework for understanding how systemic nutritional changes selectively exacerbate inflammation in metabolically responsive barrier tissues such as the skin.

In summary, our study defines the FABP5/MAPK/CCL20 axis in keratinocytes as a new link between saturated fat and psoriatic inflammation. By coupling saturated fatty acids to MAPK/CCL20 signaling, FABP5 enables metabolically driven crosstalk between keratinocyte and IL-17-producing γ δ T cells, thereby amplifying local psoriatic inflammation. Our study provides a mechanistic framework for understanding how obesity and dietary lipids exacerbate psoriasis at specific tissue sites and highlights FABP5 as a potential therapeutic target for obesity-driven psoriatic inflammation.

## Author contributions

J.Y. performed the experiments and analyzed the data. J.H., A.A., X.J., and X.H. provided technical assistance. B.L. designed the experiments. A.J. helped with intellectual inputs and paper writing. J.Y., and B.L. wrote the paper.

## Data Availability Statement

All data generated in this study will be made available upon request from the lead contact. This paper does not report original code. Any additional information required to reanalyze the data reported in this paper is available from the lead contact upon request.

## Abbreviations

CCL20: C-C motif chemokine ligand 20;
FABP5: fatty acid-binding protein 5;
HFD: high-fat diet;
MAPK: mitogen-activated protein kinase;
SFA: saturated fatty acid;
TCR: T-cell receptor.

