## Supplemental materials for "Keratinocyte FABP5 mediates saturated fat-induced MAPK/CCL20 signaling in psoriatic inflammation"

###### **Supplemental Figure 1-5**

###### **Supplemental Table S1**

Figure-S1

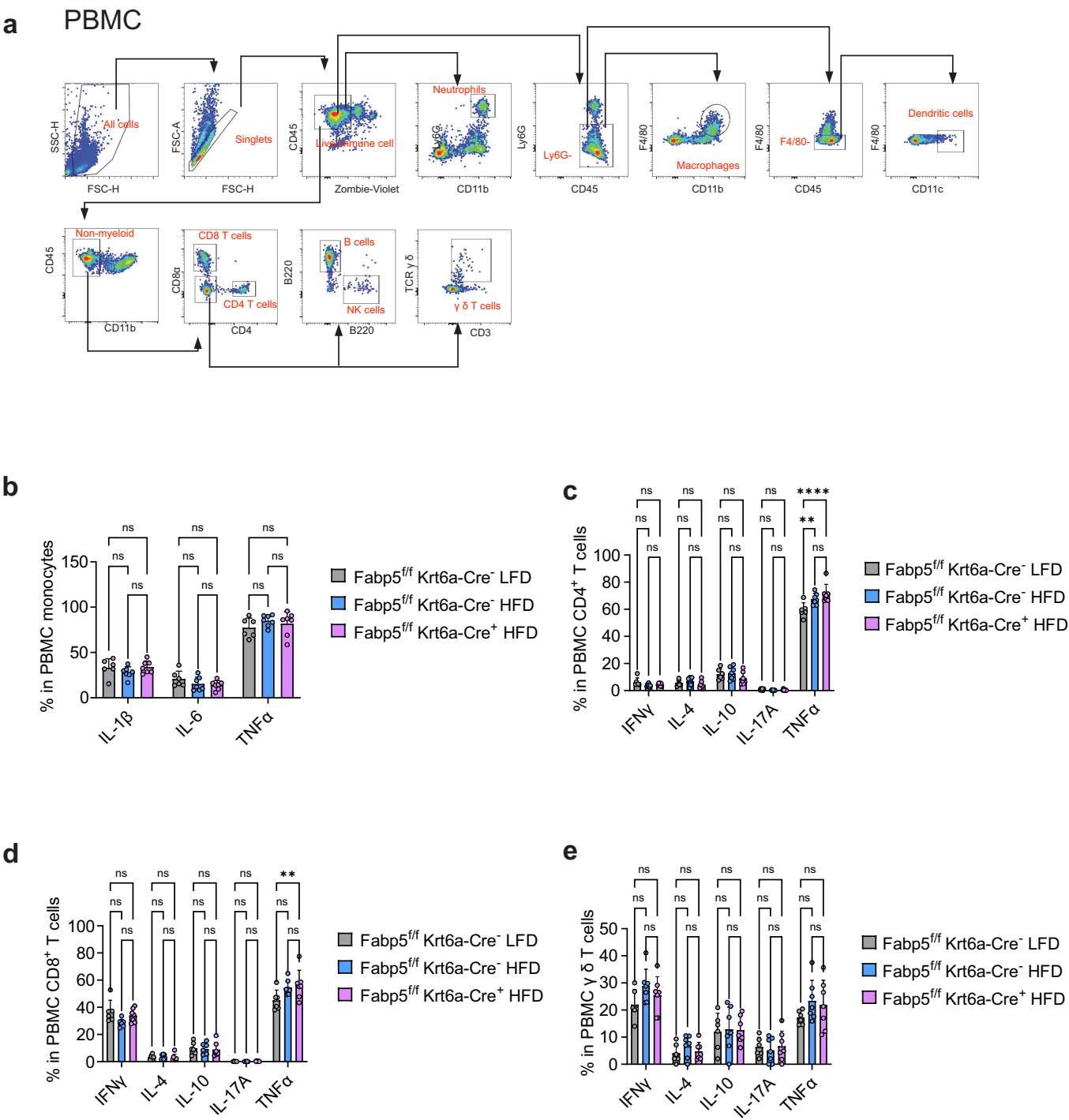

**Supplemental Figure 1. Baseline cytokine production in PBMC subsets from Fabp5<sup>ff</sup> Krt6a-Cre mice fed saturated HFD or LFD.**

(a) Proposed gating strategy for mouse PBMC subsets. (b-e) Cytokine production by flow cytometry in monocytes (b), CD4<sup>+</sup> T cells (c), CD8<sup>+</sup> T cells (d), and $\gamma$ $\delta$ T cells (e) of PBMCs from Fabp5<sup>ff</sup> Krt6a-Cre mice fed saturated HFD or LFD for 5 months. Data are shown as mean ± SD, statistical significance was determined using two-way ANOVA with Tukey's multiple comparison test. \*\*, $P < 0.01$; \*\*\*, $P < 0.001$; ns, nonsignificant.

### Figure-S2

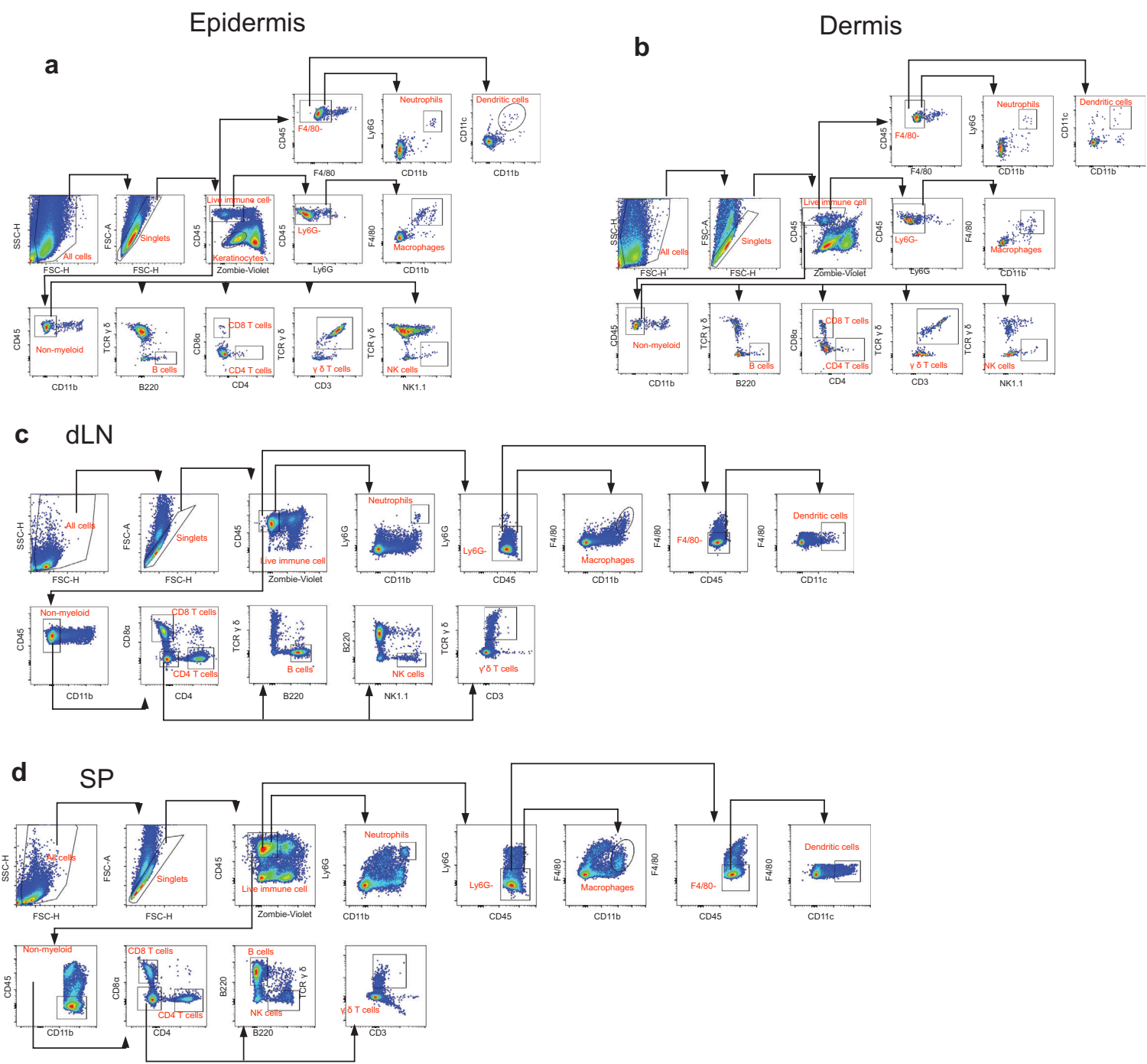

**Supplemental Figure 2. Flow cytometric gating strategies for multi-organ immune subsets in saturated HFD-related psoriasis.**

(a-d) Proposed gating strategy for immune subsets in epidermis (a), dermis (b) draining lymph nodes (dLN) (c), and spleen (d) in *Fabp5<sup>ff</sup>* Krt6a-Cre mice fed saturated HFD or LFD for 5 months, followed by 4-days IMQ treatment.

### Figure-S3

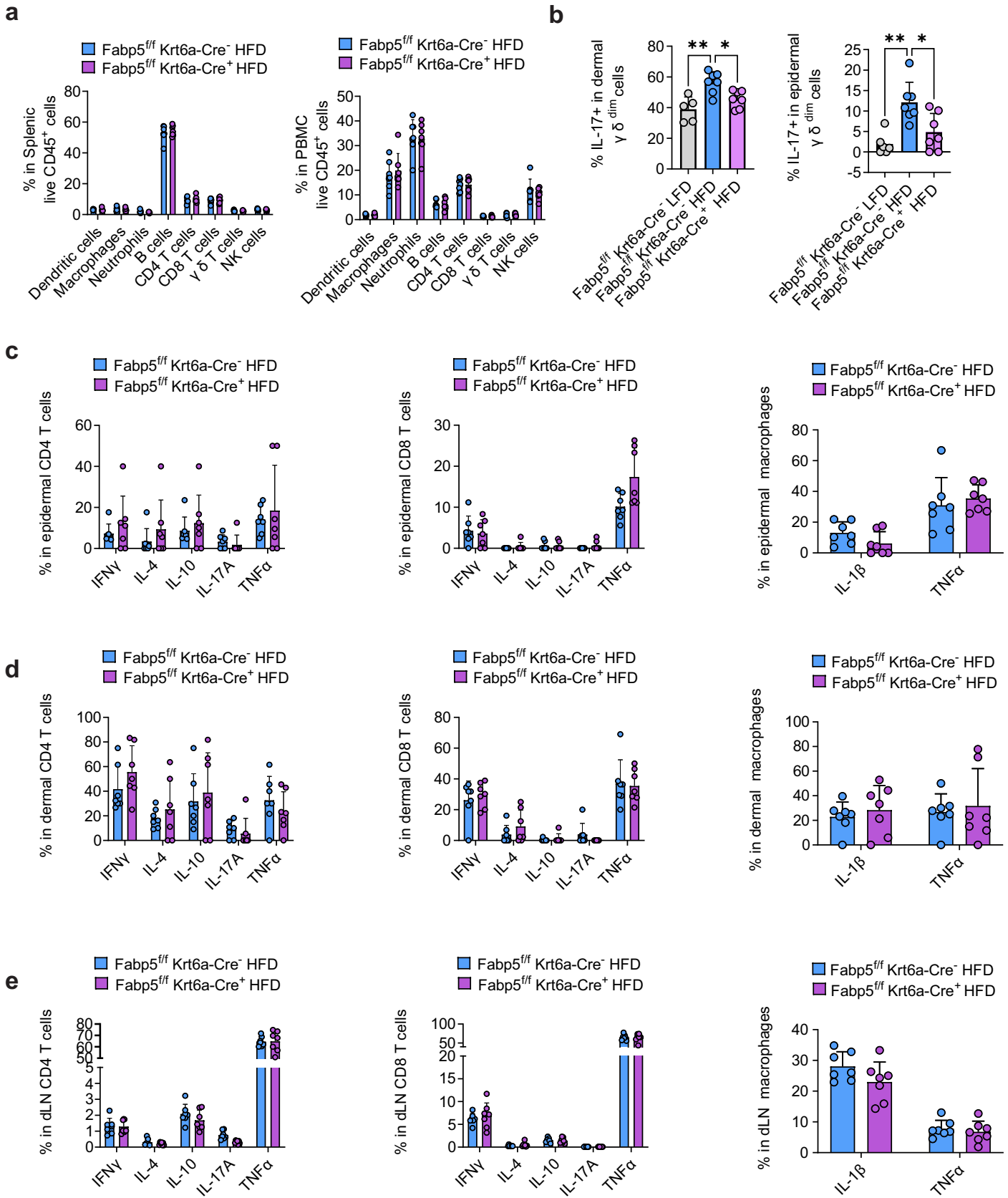

**Supplemental Figure 3. Immunophenotypic flow staining of Fabp5<sup>ff</sup> Krt6a-Cre mice in saturated HFD-related psoriasis.**

(a) Immunophenotype in spleen and PBMC of Fabp5<sup>ff</sup> Krt6a-Cre mice fed saturated HFD or LFD for 5 months, followed by 4-days IMQ treatment. (b) Statistical analysis of IL-17A+ ratio in TCR $\gamma \delta^{\text{dim}}$ cells in epidermis and dermis from Fabp5<sup>ff</sup> Krt6a-Cre mice fed saturated HFD or the LFD for 5 months, followed by 4 days of IMQ treatment. (c-e) Cytokine production by flow cytometry in CD4<sup>+</sup> T cells, CD8<sup>+</sup> T cells, and macrophages in epidermis (c), dermis (d), and dLNs (e) from Fabp5<sup>ff</sup> Krt6a-Cre mice fed saturated HFD or LFD for 5 months, followed by 4-days IMQ treatment. Data are shown as mean ± SD, statistical significance was determined using one-way ANOVA with Bonferroni's multiple comparison test for (b) and two-way ANOVA with Tukey's multiple comparison test for (a, c-e). \*, P < 0.05; \*\*, P < 0.01.

### Figure-S4

a

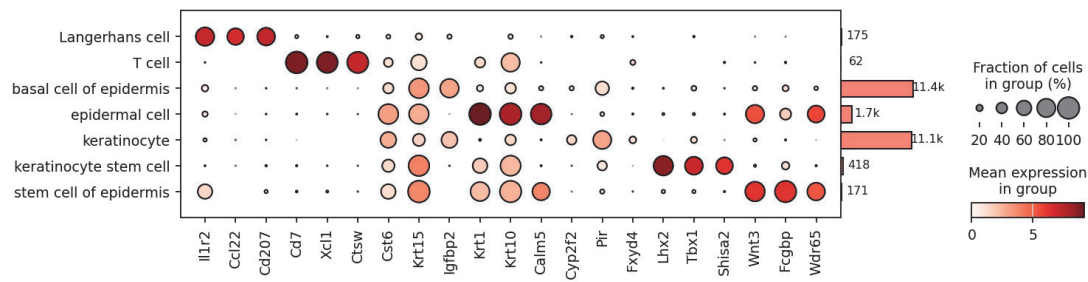

b

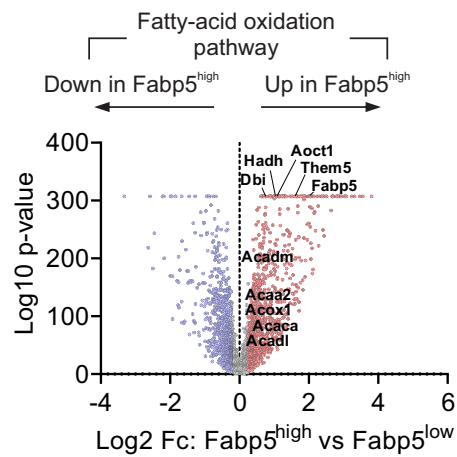

c

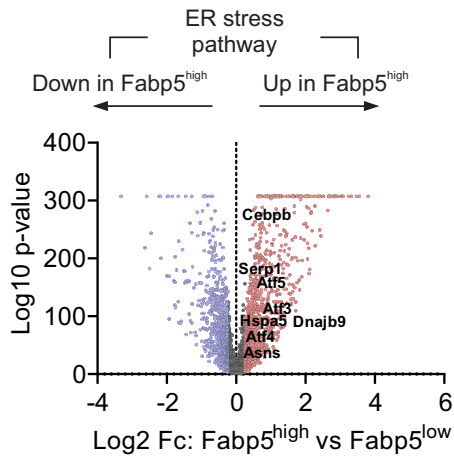

d

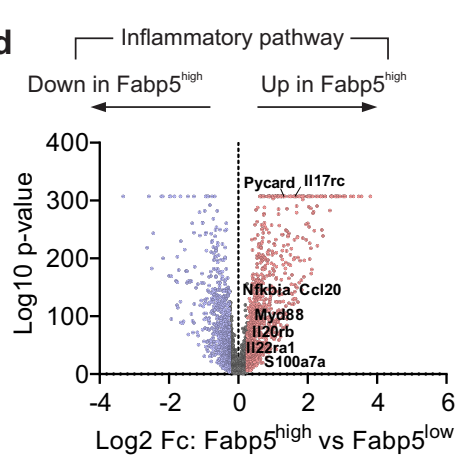

e

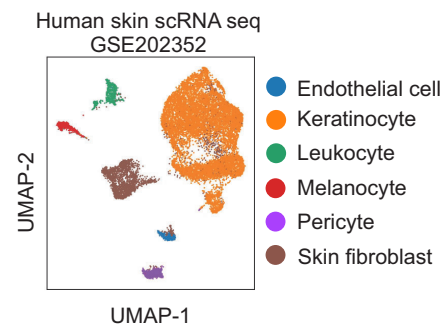

f

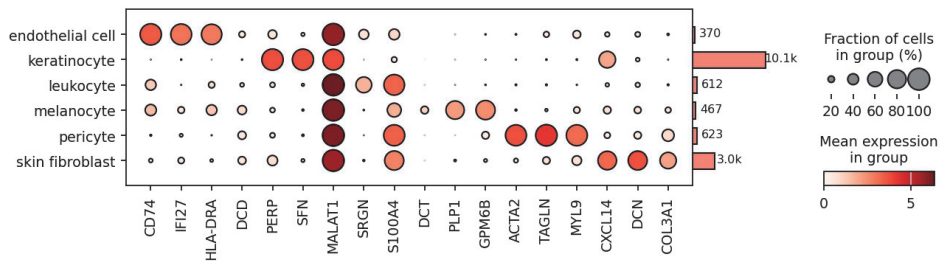

g

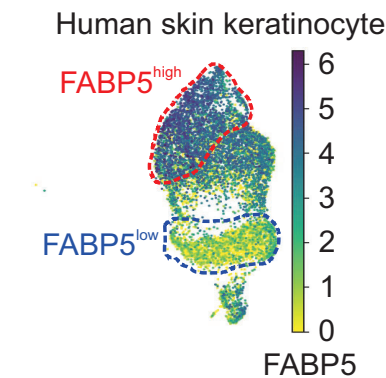

h

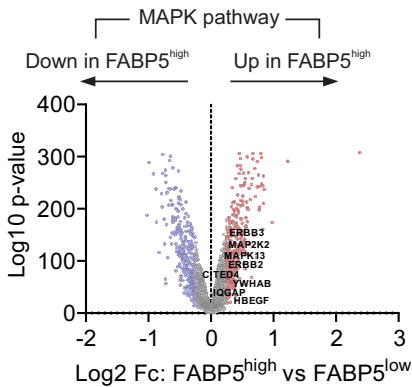

i

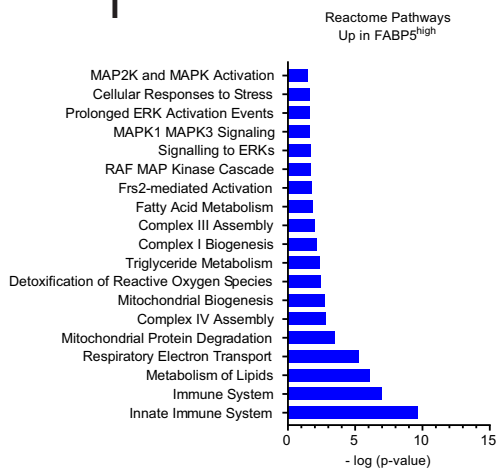

**Supplemental Figure 4. Single-cell RNA sequencing (scRNA-seq) analysis of FABP5<sup>high</sup> and FABP5<sup>low</sup> skin keratinocytes in mice and humans.**

(a) Dot plot of canonical cell lineage-defining markers for the populations annotated in Figure 5a. DEG analysis of genes involved in MAPK pathways (b), ER stress pathways (c), and Inflammatory pathways (d), comparing Fabp5<sup>high</sup> and Fabp5<sup>low</sup> mouse keratinocytes shown in volcano plot with an adjusted p value cutoff of 0.05 and a Log2 fold change cutoff of $\pm 0.26$. Red dots are significantly upregulated genes in Fabp5<sup>high</sup>, blue dots are significantly downregulated genes in Fabp5<sup>high</sup>, and gray dots are genes with no significant difference. (e) ScRNA-seq clustering analysis of human skin tissue. (f) Dot plot of canonical cell lineage-defining markers for the populations annotated in (e). (g) Identification of FABP5<sup>high</sup> and FABP5<sup>low</sup> human keratinocytes based on FABP5 gene expression level in ScRNA-seq. (h) DEG analysis of genes involved in MAPK pathways comparing FABP5<sup>high</sup> and FABP5<sup>low</sup> human keratinocytes shown in volcano plot with an adjusted p value cutoff of 0.05 and a Log2 fold change cutoff of $\pm 0.26$. Red dots are significantly upregulated genes in Fabp5<sup>high</sup>, blue dots are significantly downregulated genes in Fabp5<sup>high</sup>, and gray dots are genes with no significant difference. (i) Pathway enrichment analysis of significantly upregulated genes in FABP5<sup>high</sup> human keratinocytes using Reactome Pathways database.

Figure-S5

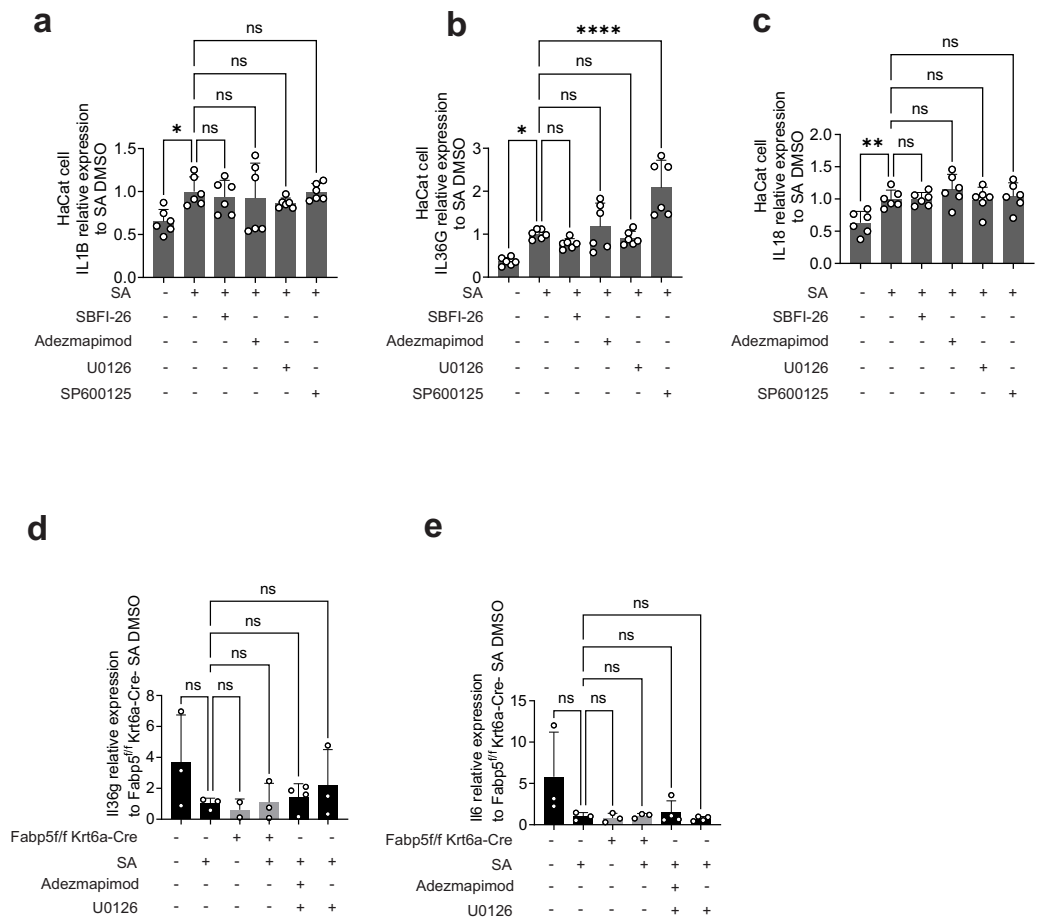

**Supplemental Figure 5. Cytokine expression levels in HaCat cells and mouse primary keratinocytes by qPCR.**

(a-c) Expression levels of IL1B (a), IL36G (b), IL18 (c) by qPCR in HaCat pretreated different inhibitors for 1-hours followed by 8 hours of fatty-acid treatment in the continued presence of inhibitors. (d-e) Expression levels of Il36g (d) and Il6 (e) by qPCR in the primary epidermal cells isolated from Fabp5<sup>ff</sup> Krt6a-Cre mice. Data are shown as mean ± SD, statistical significance was determined using one-way ANOVA with Bonferroni's multiple comparison test. \*, $P < 0.05$; \*\*, $P < 0.01$; \*\*\*\*, $P < 0.0001$; ns, no significance.

**TABLE S1 Real-time PCR primers**

| **Name** | **Forward Sequence (5'-3')** | **Reverse Sequence (5'-3')** |
| --- | --- | --- |
| **mFabp5** | AACCGAGAGCACAGTGAAG | ACACTCCACGATCATCTTCC |
| **mCcl20** | AAGACAGATGGCCGATGAAG | AGTCCACTGGGACACAAATC |
| **mIl23a** | CAGAACCAAAGGAGGTGGATAG | GAAGGATCTTGGAACGGAGAAG |
| **mIl36g** | GAGAGAGCTGGGCTATTTGTATC | ACATCCTTACACCACTCCAATC |
| **mIl6** | CTTCCATCCAGTTGCCTTCT | CCTTCTGTGACTCCAGCTTATC |
| **mHprt1** | AGCCTAAGATGAGCGCAAGT | TTACTAGGCAGATGGCCACA |
| **hCCL20** | TTGATGTCAGTGCTGCTACTC | TCTTGGGCTATGTCCAATTCC |
| **hIL1B** | CAAAGGCGGCCAGGATATAA | AAGTGAGTAGGAGAGGTGAGAG |
| **hIL36G** | GGGCCGTCTATCAATCAATGTG | CTGTCACTTCGTGGAACTGC |
| **hIL18** | GCTTCCTCTCGCAACAAACTA | CATACCTCTAGGCTGGCTATCT |
| **hGAPDH** | GAAGACGGGCGGAGAGAAAC | GCCCAATACGACCAAATCCGT |
